# Species Distribution of *Cannabis sativa*: Past, Present and future

**DOI:** 10.1101/2024.06.11.598429

**Authors:** Anna Halpin-McCormick, Tai McClellan Maaz, Michael B. Kantar, Kasey E. Barton, Rishi R. Masalia, Nick Batora, Kerin Law, Eleanor J. Kuntz

## Abstract

*Cannabis sativa* L. is an annual flowering herb of Eurasian origin that has long been associated with humans. Domesticated independently at multiple locations at different times for different purposes (food, fiber, and medicine), these long-standing human associations have influenced its distribution. However, changing environmental conditions and climatic fluctuations have also contributed to the distribution of the species and define where it is optimally cultivated. Here we explore the shifts in distribution that *C. sativa* may have experienced in the past and explore the likely shifts in the future. Modeling under paleoclimatic scenarios shows niche expansion and contraction in Eurasia through the timepoints examined. Temperature and precipitation variables and soil variable data were combined for species distribution modeling in the present day and showed high and improved predictive ability together as opposed to when examined in isolation. The five most important variables explaining ∼65% of the total variation were soil organic carbon content (ORCDRC), pH index measured in water solution (PHIHOX), annual mean temperature (BIO-1), mean temperature of the coldest quarter (BIO-11) and soil organic carbon density (OCDENS) (AUC = 0.934). Climate model projections where efforts are made to curb emissions (RCP45/SSP245) and the business as usual (RCP85/SSP585) models were evaluated. Under projected future climate scenarios, shifts worldwide are predicted with a loss of ∼43% in suitability areas with scores above 0.4 observed by 2050 and continued but reduced rates of loss by 2070. Changes in habitat range have large implications for the conservation of wild relatives as well as for the cultivation of *Cannabis* as the industry moves toward outdoor cultivation practices.

## Introduction

*Cannabis sativa* is an annual diecious herb of Eurasian origin and inhabits a range of distinct geographies and climates (Small, 2015; Long et al., 2017; McPartland & Small, 2020). These environments are characterized by having a good water supply and range of soils that tend to be well-drained, nitrogen-rich, loamy, and alluvial (Clare and Merlin, 2013; Pollio, 2016). Humans have long had a relationship with *Cannabis* with the first observations of seeds associated with pottery fragments dated to ∼10,000 years ago (Fleming and Clarke, 1998; Okazaki et al., 2011; Clarke and Merlin, 2013). This long human use has made the taxonomy of the *Cannabis* genus a major question. Historically, it has been broadly divided into two types, hemp- or drug-type, with early descriptions dating back to Linnaeus (1753) and Lamarck (1785) (Linnaeus, 1753; Lamark, 1783). Linnaeus described the plants from Northern Europe as *Cannabis sativa* and Lamarck described plants from India as *Cannabis indica*. At the time, morphological differences between the taxa led to the proposition of two species, however, recent work supports the rank of subspecies (McParland, 2018), with *Cannabis* now considered a monotypic genus (Small, 2017; Barcaccia et al., 2020; Kocalchuck et al., 2020), with genetic diversity occurring across a latitudinal gradient along which classic differentiating phenotypes occur (Zhang et al., 2018).

The domestication of *Cannabis* occurred for fiber, seed and cannabinoid content (Clarke and Merlin, 2013; Kuddus et al., 2013; Ren et al., 2021). The divergence of hemp and drug-type *Cannabis* ancestors from wild populations occurred ∼12,000 years ago, followed by a separation of hemp and drug-type gene pools occurring ∼4,000 years ago (Ren et al., 2021). Hemp-type *Cannabis* can be further sub-classified into Narrow Leaf Hemp (NLH) or Broad Leaf Hemp (BLH) with all hemp types classified as *C. sativa* ssp *sativa*. Pollen grains in the archeological record has revealed that NLH spread out of the putative ancestral zone in Central to Northern Eurasia moving westward across into mainland Europe around 6 million years ago (MYA) whereas in contrast BLH spread into Southern China and Southeast Asia about 1.2 MYA (McPartland et al., 2019; McPartland and Small, 2020). Similarly, drug-type *Cannabis* can be divided into Narrow Leaf Drug (NLD) originating from South Asia (named “*sativa*” in the recreational market) and Broad Leaf Drug (BLD) originating from Central Asia (referred to as “*indica*” in the recreational market) (McPartland and Guy, 2017). The NLD varieties are typically found along the Himachal Pradesh with a range that expands into the Montagne regions of Northern India, Uttarakhand, Nepal, Sikkim, Bhutan and into the Arunachal Pradesh (McPartland and Small, 2020). On the other hand, BLD varieties are reported to have spread from Jammu and Kashmir into Pakistan and Afghanistan (McPartland and Small, 2020).

*Cannabis* occurs in a range of diverse habitats, from cold and dry climates with short growing seasons in temperate regions to warm and wet climates with longer growing seasons in the tropics (Clarke and Merlin, 2013). Since its divergence from *Humulus Lupulus* between 18.23 −25.4 MYA (Zhang et al., 2018; Jin et al., 2020), *Cannabis* has been exposed to many different environmental stressors. It has been hypothesized that following the last glacial maximum (LGM), *Cannabis* plants migrated into and persisted in refugia sites (e.g., Hengduan Mountains, Yungui Plateau, Caucasus Mountains, northern Mediterranean peninsula) until conditions were favorable once again for range expansion (Clarke and Merlin, 2013). Habitat fragmentation and isolation driven by changing climate may have aided population separation and driven adaptive divergence among sub-species.

Currently, *Cannabis* cultivation occurs in both outdoor and indoor settings with the less valuable hemp plants producing fibers and seed oils usually cultivated outdoors and the high value medicinal and recreational drug-type plants often cultivated indoors or in more controlled farming systems (Monthony et al., 2021; Simiyu et al., 2022). As outdoor cultivation becomes the standard practice species distribution maps can help define and rank the importance of critical environmental properties and identify where these suitable environments may exist around the world currently and how they are likely to be affected by changing climate in the future. Further, there has been increased scientific interest in the last decade around understanding the soil determinants important for *Cannabis* cultivation (Wengert et al., 2021). Identifying the best climatic and soil conditions for growth informs nutrient and water management and approaches such as species distribution modeling (SDM) providing a means for more informed land selection that may facilitate minimizing negative environmental externalities (Mehrabi et al., 2019). Additionally, shifting climate also requires long term planning to prepare for future changes in agricultural requirements for outdoor production. Currently, one major region for *Cannabis* cultivation is in California, in particular the Emerald Triangle. This state has become a major cultivation center in recent history and continues to be the largest *Cannabis* market in the world with $5.2 billion in sales in 2021 (https://www.economist.com/united-states/2022/05/14/in-california-the-worlds-largest-legal-weed-market-is-going-up-in-smoke). Therefore, the objectives of this study were twofold 1) integrate publicly available soil data and climate data, past, present, and future with wild *Cannabis* occurrence points to understand how regions of the world have changed in suitability for *Cannabis* through time and 2) specifically explore present-day and future suitability for *Cannabis* cultivation in the state of California as this state is known worldwide for its outdoor cultivation.

## Materials and Methods

Occurrence data were obtained from iNaturalist (GBIF.org - https://doi.org/10.15468/dl.d8n6hx). This dataset contained 416 occurrence points which had paired images for each occurrence point (**Table S1**). Of these, 302 were deemed as wild or escapees and growing without human intervention (**Table S2**). After removing duplicates there were 234 observations. After filtering for a longitude greater than zero, 137 observations remained and were used for SDM model construction (**Table S3; Fig. S1**). These occurrence points were used to query the datasets examined in this study which included the WorldClim 2.1 climate data (all 19 bioclim variables for temperature and precipitation, **Table S4**) as well as monthly climate data for solar radiation, wind speed, water vapor pressure and elevation. Data was downloaded at the highest available spatial resolution of 30 seconds (∼1 km^2^) (https://www.worldclim.org – (Fick and Hijmans, 2017). Soil properties were downloaded from the global soil database ISRIC World Soil (https://www.isric.org – (Hengl et al., 2017) (**Table S5**). These bioclimatic and soil data were used to create species distribution models (SDM) using the software Maxent (Version 3.4.4 – (Phillips and Dudík, 2008) in RStudio (Version 2022.2.0.443 – (R core team, 2023). Paleoclimate data were sourced from paleoclim.org with the highest spatial resolution of 2.5 arc-minutes (∼5km) downloaded (**Table S6** – (Fordham et al., 2917) with suitability maps were created using the Maxent software (Version 3.3.4). Future climate data were sourced from the WorldClim repository for SSP245 (mitigation) and SSP585 (business as usual) for 2050 (averages for 2041-2060) and 2070 (averages for 2061-2080) at a spatial resolution of 30-arc seconds. Future climate projections were based on the Intergovernmental Panel on Climate Change (IPCC) sixth assessment report (AR6) and used shared socioeconomic pathways (SSPs) for different global climate models. Again, suitability maps were created using the Maxent software (Version 3.3.4). California soil orders were downloaded from https://databasin.org. Map overlays were created using the ‘raster’, ‘rworldmap’, ‘ggplot2’, ‘sf’ and ‘mapdata’ packages in RStudio (R core team, 2023;). Suitability maps were overlaid for the present day (1970-2000), 2050 and 2070, with a suitability cutoff score of 0.2. Acceptable suitability is defined as 0.2 for cultivated regions (Evans et al., 2010) and 0.4 for natural areas (Radomski et al., 2022). Model quality was explored using area under the curve (AUC) and the standard deviation of the AUC across replicates (SDAUC). A good model required an AUC ≥ 0.7 and SDAUC < 0.15. Shape files used for cropping raster extents included the World Administrative Boundaries, Countries and Territories shape file (https://public.opendatasoft.com/explore/dataset/world-administrative-boundaries/export/?flg=en-us&location=2,42.07882,0.00845&basemap=jawg.light), United States shapefiles (states_21basic), Asia and Russia shapefile (https://earthworks.stanford.edu/catalog/stanford-yg089df0008) and a Europe and Southwest Asia shape file (https://earthworks.stanford.edu/catalog/stanford-yf665vp7551). For assessing habitat reduction the ‘rasterio; and ‘numpy’ packages were used in PyCharm (version 2023.3.3) to identify pixels above the threshold examined (above 0.4 suitability). These files were then imported into RStudio where pixel counts were converted to km^2^ accounting for the curvature of the earth over the range of latitudes in each file. All code is available at https://github.com/ahmccormick and high resolution figures are available at https://figshare.com/authors/Anna_H_McCormick/17741367.

## Results and Discussion

### Historic changes in the distribution of *Cannabis*

Eurasia is the putative ancestral zone for *Cannabis* and thus was the focus for the historical timepoints (**Table S6, Fig. S11**). Across this geography there were many changes in the extent of suitable native habitat over geologic time. At 3.3 million years ago (mya) high suitability was observed in west and central Eurasia, as well as temperate South and East Asia (**Fig. S11A**). At 3.2 million years ago during the Mid-Pliocene Warm Period (mPWP), high suitability remained in West-central Eurasia (**Fig. S11B**), as well as temperate South and East Asia. There is also an increase in suitability in the Tibetan Plateau region (**Fig. S11B**) as well as in Bangladesh and Northern Myanmar. The mPWP offers an opportunity to examine how a warmer than present world may have affected species distribution, as climate models estimates for this time show that the mean surface temperature globally was between 2.7-4.0 °C higher than present along with high atmospheric CO_2_ concentrations (350-450 ppm compared to 280 ppm pre-industrial revolution) (Haywood et al., 2013). Highlighting the significance of the increase in global surface temperature during the mPWP, annual temperature explained nearly 60% of the variation during the mPWP (**Fig. S12B**). Between 3.2 million and 787,000-years ago there was a loss of lower suitability ranges (0.2 - 0.4) along the Tibetan Plateau. Habitat changes likely facilitated separation of Eurasian Steppe populations from those in China and the Himalayan Mountains (**Fig. S11B and S11C**). Such habitat fragmentation likely contributed to adaptive divergence, however, throughout the entire time series there remain areas between east and west Eurasia where suitability was likely sufficient (> 0.4) to maintain gene flow, preventing full speciation.

At 787,000 years before present (MIS19) there is a loss of broad suitability in the Tibetan Plateau region, however, western Eurasia, north east China, northern India, Nepal and North and South Korea maintain high suitability (**Fig. S11C**). This same patter occurs 130,000 years before present (**Fig. S11D**). At the height of the Last Glacial Maximum, 21,000 years ago the outline of the descending ice sheet is visible in the Northern latitudes, with a loss of suitability observed in the more northerly Eurasian Steppe regions (**Fig. S11E**). This ice age event also coincides with a reduction in high suitability (>0.7) in north east China. Previous work suggested that *Cannabis* populations may have been driven into glacial refugium in the Caucasus Mountain region in Europe and east of the Himalayan foothills in the Hengduan Mountain region (Clarke and Merlin, 2013). The Hengduan mountains of southwest China is a biodiversity hotspot (Kovalchuk et al., 2020) and had a high suitability during this period likely supporting refugial *Cannabis* populations during the Last Glacial Maximum (LGM) (Clarke and Merlin, 2013). The Himalayan Mountain system also remained suitable; however, it did exhibit a reduction in the more western regions of the Himachal Pradesh (**Fig. S11E**). Between 17,000 – 300 years ago (**Fig. S11F-K**), suitability did not change substantially, likely due to the shorter length of the time steps. Despite the relative stability there was an overall expansion in East Asia during the Heinrich-Stadial (**Fig. S11F**) and contraction during the Bolling-Allerod (**Fig. 1G**). This pattern of expansion in East Asia continued after the Younger Dryas Stadial (12,900-11,700) (**Fig. S11H**) and was visible from the Early-Holocene (**Fig. S11I**), Mid-Holocene (8,326 – 4,200) (**Fig. S11J**), the late Holocene (**Fig. S11K**) and into the Anthropocene (**Fig. S11L**). Variable contribution and AUCs for each era can be found in **Fig. S12**.

**Figure 1.**
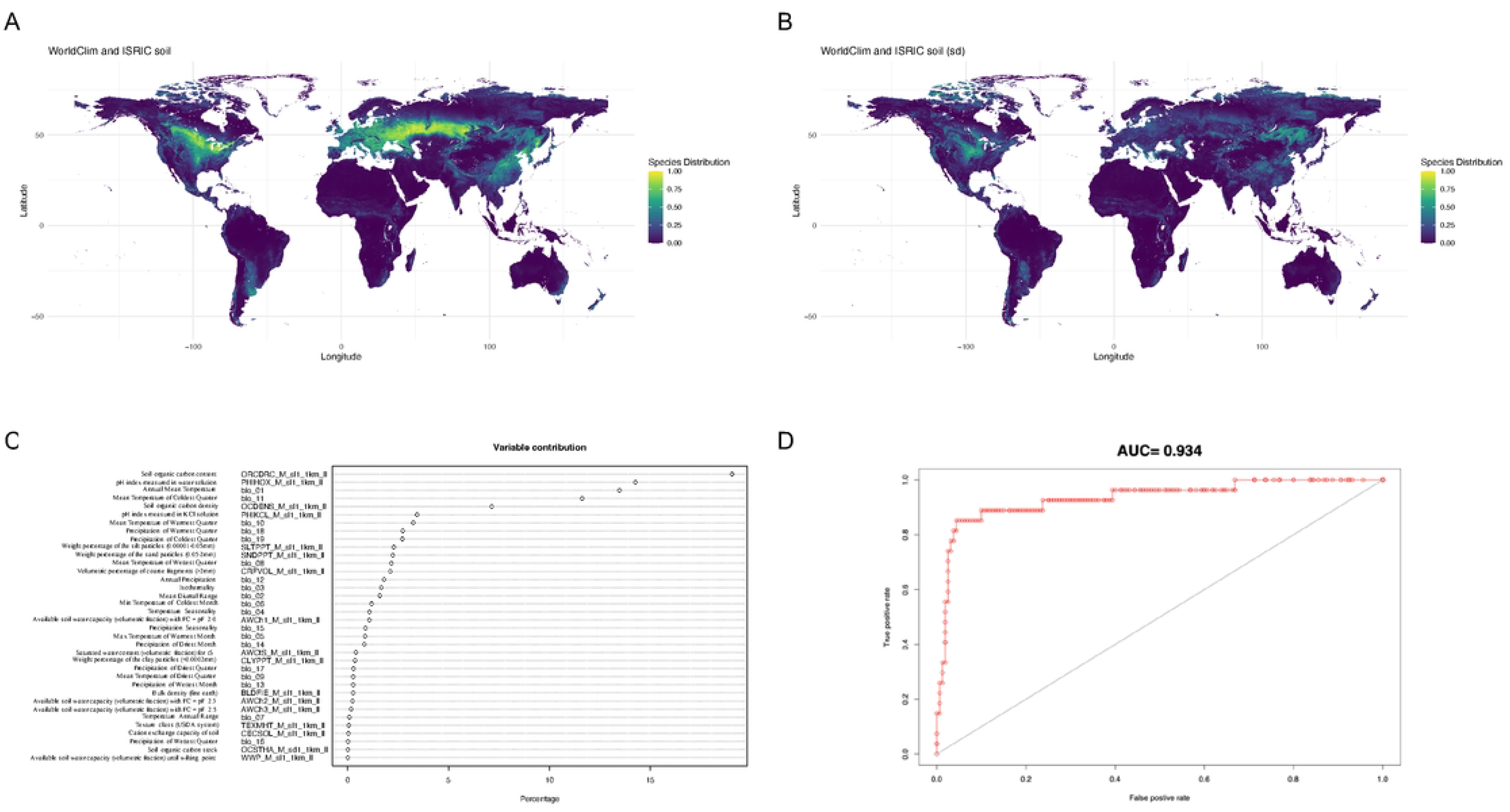
Maxent generated mean of WorldClim Bioclimatic (temperature and precipitation) variables and ISRIC soil variables for the present day together **(A)** Maxent generated suitability map for *Cannabis* cultivation worldwide for the present day using WorldClim version 2 climatic and ISRIC soil variable data compiled **(B)** Standard deviation for the merge of the WorldClim and ISRIC SoilGrid variables together. **(C)** Variable contribution of the WorldClim and ISRIC SoilGrid variables together (D) AUC for the model

Different eras showed different types of range fluctuations, for example East Asia in particular shows large suitability fluctuations during the LGM (**Fig. S11E**) as compared to the Heinrich Stadial (**Fig. S11F**), Bolling-Allerod (**Fig. S11G**) and Younger Dryas Stadial (**Fig. S11H**). High suitability is consistently observed in the Eurasian Steppe region and while the range of suitability changes throughout the Holocene (**Fig. S11I-K**), it is maintained through all the timepoints examined here (**Fig. S11A-K**). This is similarly the case for north east China and the Himalayan Mountain system (**Fig. S11A-K**). It is therefore possible that *Cannabis* may have had access to a much wider habitat range during this time than previously thought and perhaps during the LMG (**Fig. S11E**). The models here also correspond well to the subfossil pollen records which converge at the northeastern Tibetan Plateau as a proposed center of origin (McPartland et al., 2019). From here *Cannabis* it is thought to have first dispersed west to Europe by 6 MYA and to eastern China by 1.2 MYA (McPartland et al., 2019). The development of agricultural practices and the establishment of trade routes along the Eurasian Steppe (e.g. Silk Road) likely facilitated range expansion. Archeological evidence covering this expanse is still needed to support these findings.

### Current Habitat Suitability of *Cannabis*

Distributions were constructed using current (1970—2000) bioclimatic (temperature and precipitation) and soil properties separately (**Fig. S2A-B, S3A-B, S4A-B**) and together (**Fig. 1**) to explore global suitability. Six temperature and precipitation variables (BIO-1, BIO-11, BIO-10, BIO-18, BIO-19, BIO-14 - see **Table S4** for definitions) explained ∼81% of the total variation (**Fig. S3A**), with an AUC of 0.9 (**Fig. S4A**). When exploring bioclimatic variables alone, highest suitability was found in mixed deciduous forest, temperate forest steppe and taiga of Eurasia and North America (**Fig. S2A**). When exploring soil alone, four variables (ORCDRC, PHIHOX, OCDENS, CECSOL - see **Table S5** for definitions) explained ∼79% of the total variation (**Fig. S3B**) with an AUC of 0.939 (**Fig. S4B**). Here suitability was also found in the mixed deciduous forest, temperate forest steppe and taiga of Eurasia and North America (**Fig. 2B**).

**Figure 2.**
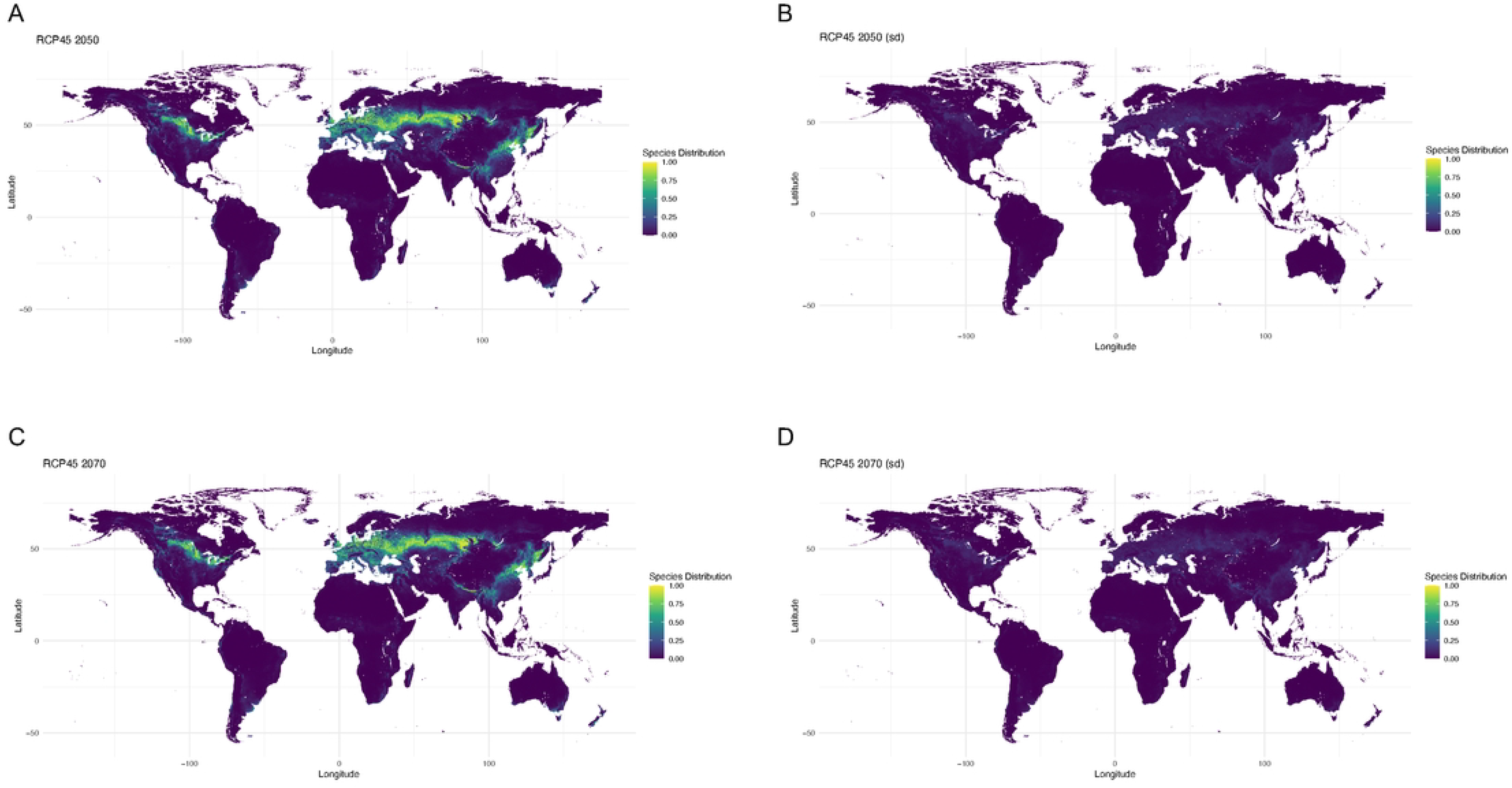
Future climate projections for 2050 and 2070 with model SSP245 (efforts made to curb climate change) **(A)** Worldwide suitability map for 2050 **(B)** Standard deviation for the worldwide suitability map for 2050 **(C)** Worldwide suitability map for 2070 **(D)** Standard deviation for the worldwide suitability map for 2070.

When climate and soil variables were combined (**Fig. 1**) the most important variables were soil organic carbon content (ORCDRC), pH index measured in water solution (PHIHOX), annual mean temperature (BIO-1), mean temperature of the coldest quarter (BIO-11) and soil organic carbon density (OCDENS) (**Fig. 1C**). With an AUC of 0.934 these five variables explained ∼65% of the total variation (**Fig. 1D**). Integrating these two datasets we see a notable improvement in model performance (**Fig. 1A**), specifically in mitigating the occurrence of anomalous suitable areas, for example those previously identified at high latitudes of Northern Canada for the ISRIC soil dataset when examined in isolation (**Fig. S2B**). *Cannabis* is thought to have had a Central Asian Origin (de Candolle, 1885; McPartland, 2018) or an origin region spanning Eurasia (Herder, 1892; Vavilov, 1926). Here, high broad suitability (above 0.4) is seen in these regions with a range spanning Europe (**Fig. S6**), Russia and Asia (**Fig. 1A, 4A, S6**). The patterns of suitability observed matches present-day reports of extant *Cannabis* populations (Clarke and Merlin, 2013). High suitability is also seen in other continents such as regions of North America (**Fig. 1A, 4A, S9**).

To examine how other environmental variables may affect the model, additional environmental properties namely; solar radiation (kJm -2/day) (**Fig. S2C**), wind speed (m/s) (**Fig. S2D**), water vapor pressure (kPa) (**Fig. S2E**), and elevation (m) (**Fig. S2F**) were acquired. Bringing together all of these environmental properties, a similar pattern of suitability is observed across Eurasia and North America. However, there is a more pronounced variability in the standard deviation for the model worldwide (**Fig. S5B**) in contrast to the more localized variations seen in the bioclimatic and soil standard deviations (**Fig. 1B**). Examining variable contribution across all 73 layers (**Fig. S6A**) shows that 6 of the top 10 contributors are bioclimatic variables (BIO-1, BIO-9 and BIO-11) and soil (ORCDRC, PHIHOX, PHIKCL) properties (**Fig. 6A**) and suggests that integrating bioclimatic and soil properties may reduce background errors that could arise when a broader array of variables is included in the modeling process and which have low variable contributions.

In the important production area of California, good suitability occurs in many counties including Sonoma, Napa, Marin County, San Meteo, Almeda, Monterey, Suter, Yuba and Butte, San Luis Obispo, Santa Barbara, Placer, Mendocino and Humboldt County and almost all coastal counties (**Fig. 4C, S10A**). Additional states which are known for outdoor cultivation were also examined for suitability for all six environmental properties examined namely; Colorado (**Fig. S10B**), Maine (**Fig. S10C**), Oregon (**Fig. S10D)**, Washington (**Fig. S10E**), Massachusetts (**Fig. S10F**) and Michigan (**Fig. S10G**).

### Suitability and Soil

Like many plant species, making use of soil data increases distribution model performance (Kantar et al., 2015). Important soil determinants for *Cannabis* suitability included soil pH, soil organic carbon, and cation exchange capacity (CEC) (**Fig. S3B**), which together regulate soil fertility and nutrient cycling in soil. Others have previously reported that *Cannabis* favors growing in alluvial soils under slightly acidic conditions in temperate and subtropical regions (Clarke and Merlin, 2013). Alluvial soils can be rich in organic matter and plant nutrients, and thus relatively fertile (Boettinger, 2005), and such soils are found along the Himachal Pradesh (Shirur et al., 2017) and in many places within the global model that have good suitability for *Cannabis* cultivation. Taxonomically, alluvial soils are diverse and fall under an assortment of soil classes (Boettinger, 2005). Likewise, in the case of *Cannabis*, higher suitability is observed in regions defined by an array of classifications, including the soil orders of Mollisols (Central Eurasian Steppe and North America), Inceptisols and Ultisols (both found predominantly in southern Russia, Mongolia, and China), with some smaller suitable regions found in areas consisting of Alfisols and Spodosols (Western Eurasian steppe) (**Fig. S2B**). This range in soil taxonomic classifications may highlight the plastic and generalist nature of *Cannabis*. In California, we found a correlation between *Cannabis* suitability and the soil orders of Inceptisols, Mollisols, Alfisols, Aridisols, and Ultisols (**Fig. 4C-D**). The weakly developed Inceptisols of California have a wide range of characteristics, but are mostly associated with steeply sloped chaparral or montane conifer forests and along streams (O’Geen et al., 2007). Certain Inceptisols in Northern California were also developed from volcanic deposits (Andisols) and have slightly acidic pH with highly variable cation exchange capacity. Mollisols define a large proportion of the suitability score that we see in California (**Fig. 4C-D**), North America (**Fig. 2B**), and the Eurasian Steppe (**Fig. S2B**). Mollisols are inherently fertile with over 50% base saturation, abundant organic matter, and near neutral pH; whereas moderately weathered Alfisols have more than 35% base saturation with clay rich subsoil horizons with high cation exchange capacity having undergone less intensive leaching than Ultisols (O’Geen et al., 2007; NRCS, 2022).

### Suitability of *Cannabis under Future Climate Scenarios*

Future distributions were constructed for both 2050 and 2070 under two emission scenarios (SSP245-curbing emissions and SSP585 – business as usual). Under SSP245 there is decreasing suitability by 2050 (**Fig. 2A**) and 2070 (**Fig. 2C**). Under SSP585 there is an even greater decrease in suitability by 2050 (**Fig. 3A**) and 2070 (**Fig. 3B**). Future climate datasets represent climatic variables only, as soil properties are more challenging to estimate. Despite the loss of suitability in the future there was overlap between suitability ranges in 2050 and 2070 for both SSP scenarios (**Fig. 4A**).

**Figure 3.**
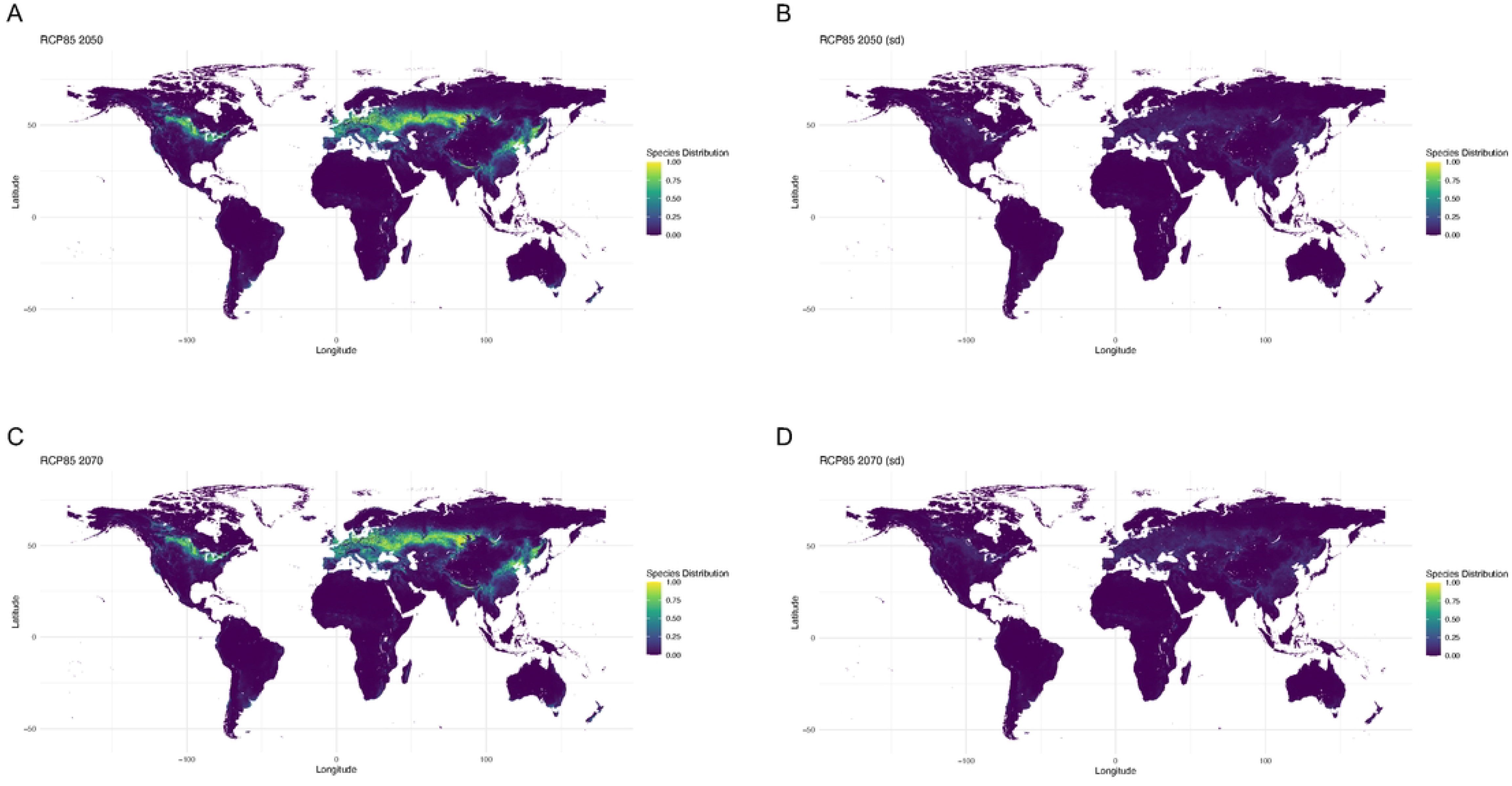
Future climate projections for 2050 and 2070 with model SSP845 (business-as-usual climate model) **(A)** Worldwide suitability map for 2050 **(B)** Standard deviation for the worldwide suitability map for 2050 **(C)** Worldwide suitability map for 2070 **(D)** Standard deviation for the worldwide suitability map for 2070.

**Figure 4.**
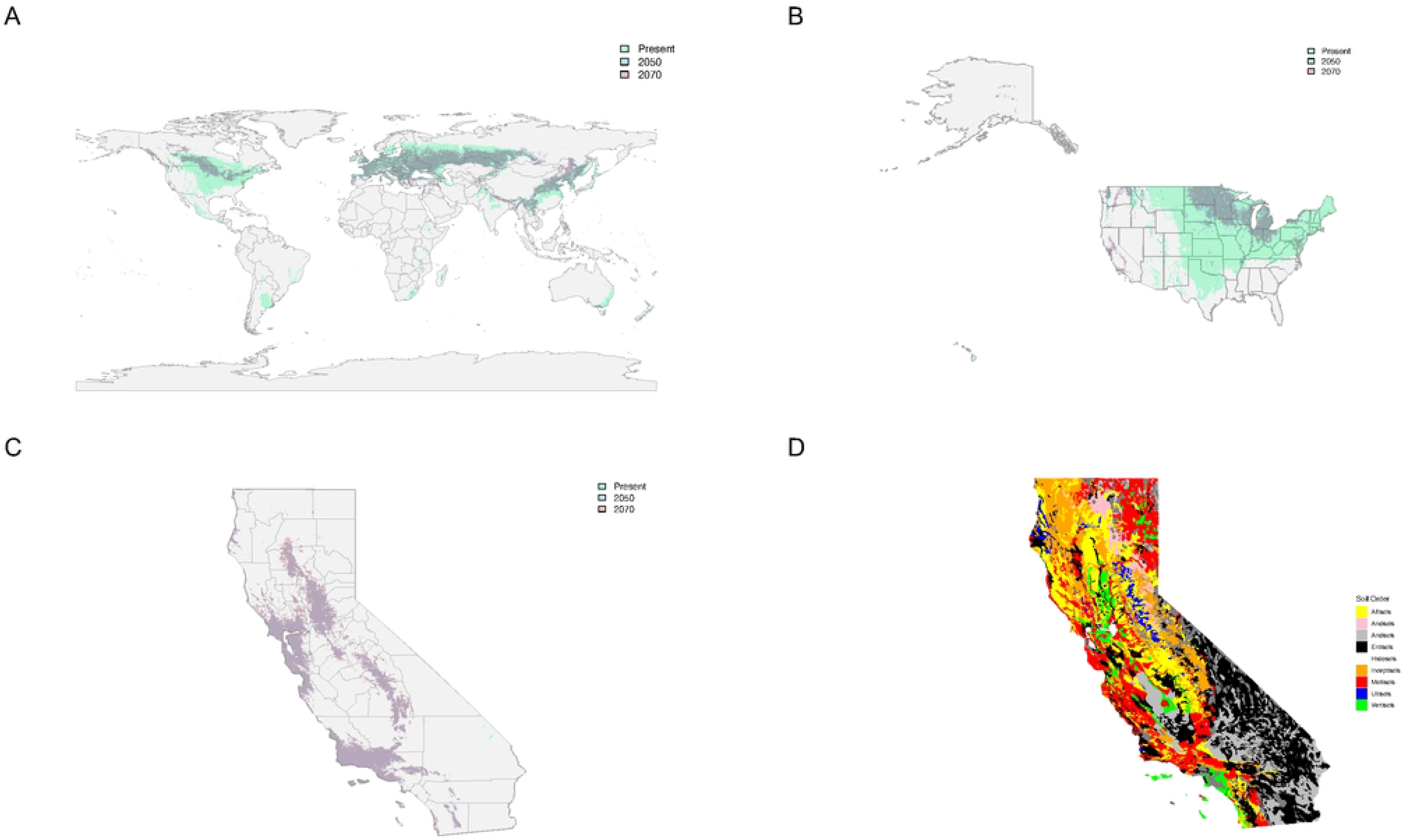
Suitability overlay with the present day (green) and future projections for *Cannabis sativa* species distribution for 2050 (blue) and 2070 (red) for SSP845 or the business-as-usual climate model **(A)** Worldwide suitability map **(B)** Suitability map for the United States and **(q** Suitability map for the state of California, where green represents present day suitability (above 0.2), Blue represents suitability (above 0.2) for 2050, red suitability (above 0.2) for 2070 and purple the overlap of suitability for 2050 and 2070 **(D)** Soil orders for the stateof California.

Partitioning by subregions such as Asia and Russia (**Fig. S7A-E**), Europe and SE Asia (**Fig. S8 A-E**) and the United States (**Fig. S9A-E**) allows for a more quantitative examination of the changes these areas may experience from the present day moving into 2050 and 2070. The rasters generated for each climate scenario (SSP45 and SSP85) and each future time point (2050 & 2070) were filtered for suitability values above 0.4, which is considered an acceptable suitability value for natural growth. Pixel counts were performed and converted into km^2^, accounting for the curvature of the earth within the range of latitudes in the partition. Worldwide across all climate models and timepoints we see an average of ∼43% reduction in suitable area (**Table S7**) with a reduction from ∼13.8 million to ∼7.8 million km^2^. Within the partition of Asia and Russia a ∼29% reduction in suitable area from ∼6.4 million to ∼4.5 million km^2^ is observed (**Fig. S7A-E, Table S7**) and similarly in Europe ∼15% losses are observed with a reduction from ∼2.5 million to ∼2.2 million km^2^ (**Fig. S8A-E, Table S7**). In the United States an average of ∼81% loss in suitable area is observed with a reduction from ∼2.8 million to ∼0.5 million km^2^ (**Fig. S9A-E, Table S7**).

Within the United States, there was a contraction in suitability (**Fig. 4B, Fig. S8A-E**), although, suitability remains favorable in areas of California (**Fig. 4C**), with many important production counties showing favorable suitability scores (e.g., Humboldt, Mendocino Sonoma, Marin, Santa Barbara, and Ventura) (**Fig. 4C**). Again, *Cannabis* suitability was correlated with areas where Inceptisols, Mollisols, Vertisols, Ultisols and Alfisols are predominantly found (**Fig. 4C-D**).

### Implications for Future Cultivation

Globally combining climate and soil variables provides improved resolution (**Fig. 1A**). The observed suitable areas (>0.4) match common agricultural lands known for *Cannabis* cultivation in Asia and Europe and captured specific geographies where genetic studies have shown the presence of drug-type feral *Cannabis* populations (Ren et al., 2021). The overlap between suitability for 2050 and 2070 suggests that the effects on *Cannabis* distribution due to changing climate will have mostly taken effect by 2050 (**Fig. 4**). Moving into the future, in North America, the states of North Dakota, Minnesota, Wisconsin, and Michigan show high suitability (**Fig. 4B**).

Areas in the states of Colorado, Wyoming and California show moderate suitability for the present day, however, moving into the future these areas show decreasing suitability (**Fig. S9**). California is known worldwide for its cultivation of *Cannabis*. Proposition 64 in 2016 brought legalization to the recreational use of *Cannabis* and saw a movement towards legal operations in the production and sales of *Cannabis*. Exploring this region more specifically we observed an increase in suitability from the present day to 2050 and 2070 in areas such as Marin County, Contra Costa, San Mateo, Santa Barbara and Ventura (**Fig. 4C**). Counties which currently produce *Cannabis* may be impacted under different climate change scenarios in the future. However, future models here only take into account abiotic factors and biotic pressures will also have significant roles in future species distribution.

## Conclusion

Global suitability for *Cannabis* was explored with the intent of identifying regions where *Cannabis* cultivation could be facilitated by suitable climate and soil properties. Across Eurasia and the United States there will be broad loss of suitability by 2050 for two different emission scenarios (SSP245 and SSP585). As suitable habitat decreases, conservation of wild relatives and naturalized populations of *Cannabis* around the world will be critical for the preservation of diversity and act as a valuable source of variation for trait improvement as the industry continues to develop.

## Acknowledgments

We would like to thank Koa the University of Hawai’i (UH) high performance computing (HPC) cluster.

## Supplemental Figure Legends

**Fig. S1**. 137 observations used for SDM model construction with a longitude greater than zero.

**Fig. S2**. Individual species distributions for each set of environmental properties examined **(A)** WorldClim Bioclimatic variables **(B)** ISRIC soil data **(C)** Solar radiation (kJm^2^/day) **(D)** Wind speed (m/s) **(E)** Water vapor pressure (kPa) **(F)** Elevation suitability maps.

**Fig. S3**. Variable contribution graphs for each set of environmental properties examined **(A)** WorldClim Bioclimatic variables **(B)** ISRIC soil data **(C)** Solar radiation (kJm^2^/day) **(D)** Wind speed (m/s) **(E)** Water vapor pressure (kPa) and **(F)** Elevation.

**Fig. S4**. Area under the curve graphs each set of environmental properties examined **(A)** WorldClim Bioclimatic variables **(B)** ISRIC soil data **(C)** Solar radiation (kJm^2^/day) **(D)** Wind speed (m/s) **(E)** Water vapor pressure (kPa) and **(F)** Elevation.

**Fig. S5**. Overlay of all six environmental datasets **(A)** Worldwide plot **(B)** standard deviation for the overlay of all six environmental variables.

**Fig. S6**. Overlay of all six environmental datasets **(A)** Variable contribution graph (**B)** Area under the curve graphs each set of environmental properties examined

**Fig. S7**. Species distribution with temperature and precipitation data in Asia and Russia for **(A)** present day **(B)** SSP45 2050 **(C)** SSP45 2070 **(D)** SSP85 2050 **(E)** SSP85 2070.

**Fig. S8**. Species distribution with temperature and precipitation data in Europe for **(A)** present day **(B)** SSP45 2050 **(C)** SSP45 2070 **(D)** SSP85 2050 **(E)** SSP85 2070.

**Fig. S9**. Species distribution with temperature and precipitation data in the United States for **(A)** present day **(B)** SSP45 2050 **(C)** SSP45 2070 **(D)** SSP85 2050 **(E)** SSP85 2070.

**Fig. S10**. Species distribution for a subset of the United States with data for all six environmental properties examined **(A)** California **(B)** Colorado **(C)** Maine **(D)** Oregon **(E)** Washington **(F)** Massachusetts **(G)** Michigan.

**Fig. S11. (A)** Pleistocene: M2 (ca. 3.3 Ma) **(B)** Predicted Distribution for the Paleoclimate timepoint of the Mid Pliocene warm period (ca. 3.2 Ma) **(C)** Predicted Distribution for the Paleoclimate timepoint of the Pleistocene: MIS19 (ca. 787,000 years ago) **(D)** Predicted Distribution for the Paleoclimate timepoint of the Pleistocene: Last Interglacial (130,000 years ago) **(E)** Predicted Distribution for the Paleoclimate timepoint of the Pleistocene: Last Glacial Maximum (ca. 21,000 years ago) **(F)** Potential Distribution for the Paleoclimate timepoint of the Pleistocene: Heinrich Stadial (14,700 – 17,000 years ago) **(G)** Potential Distribution for the Paleoclimate timepoint of the Pleistocene: Bolling-Allerod (12,900 – 14,700 years ago) **(H)** Potential Distribution for the Paleoclimate timepoint of the Pleistocene: Younger Dryas Stadial (11,700 – 12,900 years ago) **(I)** Potential Distribution for the Paleoclimate timepoint of the Pleistocene: Early Holocene, Greenlandian (8,366 - 11,700 years ago) **(J)** Potential Distribution for the Paleoclimate timepoint of the Pleistocene: Mid Holocene, Northgrippian (4,200 – 8,326 years ago) **(K)** Potential Distribution for the Paleoclimate timepoint of the Pleistocene: Late Holocene, Meghalayan (300 – 4200 years ago)

**Fig. S12**. AUC and variable contribution graphs for each timepoint from the Paleoclim dataset **(A)** Pleistocene: M2 (ca. 3.3 Ma) **(B)** Mid Pliocene warm period (ca. 3.2 Ma) **(C)** Pleistocene: MIS19 (ca. 787,000 years ago) **(D)** Pleistocene: Last Interglacial (130,000 years ago) **(E)** Last Glacial Maximum (ca. 21,000 years ago) **(F)** Heinrich Stadial (14,700 – 17,000 years ago **(G)** Bolling-Allerod (12,900 – 14,700 years ago) **(H)** Younger Dryas Stadial (11,700 – 12,900 years ago) **(I)** Early Holocene, Greenlandian (11,700-8,326 years ago) **(J)** Mid Holocene, Northgrippian (4,200 −8,326 years ago) **(K)** Late Holocene, Meghalayan (300 – 4200 years ago)

## Table Legends

**Table S1**. 416 occurrence points from iNaturalist which had paired images.

**Table S2**. Latitude and longitudes of 302 occurrence points deemed to be growing wild without human intervention.

**Table S3**. Latitude and Longitude for the 137 observations used for SDM model construction, post-filtering for a longitude greater than zero.

**Table S4**. WorldClim2 Bioclimatic variable definitions.

**Table S5**. ISRIC soil variable definitions.

**Table S6**. Paleoclimate Time Period and date range

**Table S7**. Pixel counts for present day, SSP 45 and SSP85 for 2050 and 2070 above the 0.4 threshold for the world and the partitions of Asia & Russia, Europe and SE Asia and the United States.

